# Chemoproteomics yields a selective molecular host for acetyl-CoA

**DOI:** 10.1101/2022.12.19.521087

**Authors:** Whitney K. Lieberman, Zachary A. Brown, Yihang Jing, Nya D. Evans, Isita Jhulki, Carissa Grose, Jane E. Jones, Jordan L. Meier

**Affiliations:** Chemical Biology Laboratory, Center for Cancer Research, National Cancer Institute, National Institutes of Health, Frederick, Maryland 21702, United States; Protein Expression Laboratory, Cancer Research Technology Program, Frederick National Laboratory for Cancer Research, Leidos Biomedical Research, Inc., Frederick, Maryland 21702, United States

## Abstract

Chemoproteomic profiling is a powerful approach to define the selectivity of small molecules and endogenous metabolites with the human proteome. In addition to mechanistic studies, proteome specificity profiling also has the potential to identify new scaffolds for biomolecular sensing. Here we report a chemoproteomics-inspired strategy for selective sensing of acetyl-CoA. First, we use chemoproteomic capture experiments to validate the N-terminal acetyltransferase NAA50 as a protein capable of differentiating acetyl-CoA and CoA. A Nanoluc-NAA50 fusion protein retains this specificity and can be used to generate a bioluminescence resonance energy transfer (BRET) signal in the presence of a CoA-linked fluorophore. This enables the development of a ligand displacement assay in which CoA metabolites are detected via their ability to bind the Nanoluc-NAA50 protein ‘host’ and compete binding of the CoA-linked fluorophore ‘guest.’ We demonstrate that the specificity of ligand displacement reflects the molecular recognition of the NAA50 host, while the window of dynamic sensing can be controlled by tuning the binding affinity of the CoA-linked fluorophore guest. Finally, we show the method’s specificity for acetyl-CoA can be harnessed for gain-of-signal optical detection of enzyme activity. Overall, our studies demonstrate the potential of harnessing insights from chemoproteomics for molecular sensing and provide a foundation for future applications in target engagement and selective metabolite detection.

---

Acetyl-CoA is essential for life and plays a key role in energy production and lipid metabolism.^1^ Overactivation of two enzymes responsible for human acetyl-CoA metabolism, ATP-citrate lyase (ACLY) and acetyl-CoA synthetase 2 (ACSS2), are associated with several cancers^2,3^ and have been the target of substantial therapeutic development efforts.^4,5^ Acetyl-CoA also plays an important role in epigenetic signaling, protein homeostasis, and protein synthesis due to its ability to serve as a cofactor used in the modifications of proteins^6–7^ and RNA.^7^ Over the last decade, substantial evidence indicates these processes can be regulated by subcellular acetyl-CoA levels,^8–10^ highlighting the potential for acetyl-CoA to directly link metabolism and gene expression. Emphasizing this, a recent study demonstrated that concentrations of acetyl-CoA in the nucleus are distinct from those in other organelles.^11^ These experiments relied on careful cellular fractionation in combination with liquid chromatography-mass spectrometry (LC-MS). This approach holds the advantage of being able to differentiate multiple acyl-CoA metabolites, but requires specialized equipment and has a limited capacity for measuring dynamics.

Optical sensing represents an alternative approach to study CoA metabolites.^12^ Three general classes of CoA sensors have been reported which detect malonyl-CoA,^13,14^ long-chain fatty acyl (LCFA)-CoAs,^15^ and non-thioesterified CoA.^16^ Absent from this emerging arsenal are methods to detect acetyl-CoA. The selective molecular recognition of acetyl-CoA versus CoA presents a challenge to sensor design due to the necessity to differentiate two large (>750 Da) metabolites based on replacement of a single hydrogen atom with an acetyl group that comprises only ~5% of total mass (Fig. 1a). Prior efforts at selective molecular recognition of acetyl-CoA have either failed^17^ or been limited to disruption – rather than augmentation – of acetyl-CoA binding.^16^

**Figure 1.**
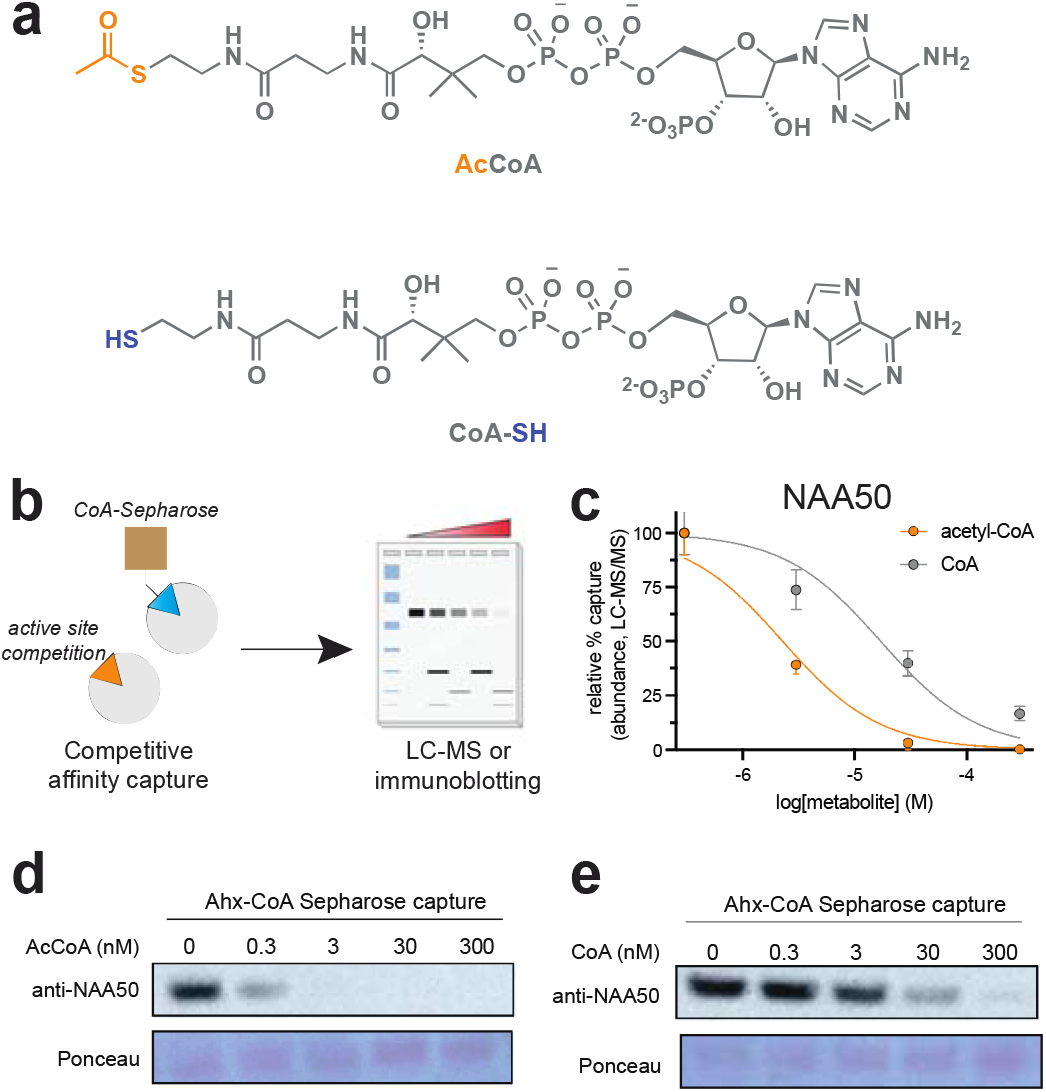
(a) Structure of acetyl-CoA and CoA. (b) Schematic of competitive chemoproteomic profiling experiment. (c) Dose-dependent competition of NAA50 capture in chemoproteomic experiments. Label-free abundance measurements were based on dNSAF, n=3. (d-e) Chemoproteomic capture of NAA50 by Ahx-CoA resins is competed by acetyl-CoA and CoA. For additional validation blotting data see Fig. S3.

Considering this challenge, we took inspiration from metabolic mechanisms of epigenetic regulation.^18,19^ Specifically, histone acetylation has been proposed to be regulated by high acetyl-CoA levels which liberate histone acetyltransferase (HAT) enzymes from metabolic feedback inhibition by CoA.^20,21^ To understand this mechanism’s selectivity our group has used chemoproteomic profiling to define how proteins bind to acetyl-CoA and CoA on a proteome-wide scale.^22^ In addition to identifying new acetyltransferases that are highly sensitive to feedback inhibition, these studies also detect enzymes that preferentially bind acetyl-CoA (Fig. 1b). To assess whether this approach could be leveraged for selective sensing, we analyzed a previous chemoproteomic dataset which determined how capture of proteins by a CoA-based affinity resin were competed by acetyl-CoA or CoA,^10^ fitting quantitative abundance data to a nonlinear regression model. Since CoA affinity resins can capture both direct and indirect interactors^23^ we focused on 23 proteins known to directly bind to acetyl-CoA,^24^ 13 of which displayed dose-dependent competition (R^2^ ≥ 0.85) by both metabolites (Fig. S1, Table S1-S2).

Comparing half-maximal inhibition (IC_50_) values, we identified three N-terminal acetyltransferases (NAA30, NAA40, NAA50) that display ≥5-fold preferential competition by acetyl-CoA (Table S2, Fig. 1c). These proteins harbor highly homologous acetyl-CoA binding sites (Fig. S2).^25^ We prioritized validation of the recognition properties of NAA50 as it yielded the strongest chemoproteomic signal (Table S2), has been well-studied from the perspective of inhibitor binding,^26,27^ and interacts with two partner proteins (NAA15 and HYPK) also selectively competed by acetyl-CoA (Table S2). Non-denatured whole cell lysates were incubated with CoA affinity resins, washed to remove non-specific interactors, and blotted for NAA50. Consistent with high-throughput measurements, acetyl-CoA competed capture of NAA50 at lower concentrations than CoA (Fig. 1d-e, Fig. S3). Surface plasmon resonance (SPR) analysis further confirmed ~10-fold preferential binding of acetyl-CoA to NAA50 (Table S3).^22,27,28^ Overall, these studies highlight the ability of chemoproteomics to characterize differential metabolite binding and specify NAA50 as a selective receptor for acetyl-CoA.

Next, we sought to exploit NAA50’s molecular recognition properties for acetyl-CoA sensing. Previous CoA biosensors have used ratiometric fluorescence^15^ and fluorescence resonance energy transfer (FRET) detection.^16^ Considering the relatively narrow sensing window of NAA50 and unknown conformational effects of acetyl-CoA binding, we wondered whether bioluminescence resonance energy transfer (BRET) – a method that can display increased detection sensitivity relative to FRET – may provide a complementary approach (Fig. 2a).^29,30^ To construct a BRET reporter for acetyl-CoA we fused NAA50 to the C-terminus of an engineered luciferase (Nanoluc/Nluc) that has shown broad utility as a BRET donor.^31^ The predicted structure the fusion protein^32^ indicated a distance of ~6 nm between the Nanoluc and NAA50 active sites, well within the 10 nm range recommended for BRET (Fig. 2b).^30^ Two CoA analogues were synthesized to serve as BRET acceptors: Cy3-Ahx-CoA (**1**) and Cy3-Ahx-D-CoA (**2**; Fig. 2c). Prior studies have found both of these molecules capture NAA50 from cell extracts when immobilized on resin; however, enrichment by Ahx-D-CoA (**2**) is more efficient than **1** due to its higher affinity.^33^ Our design rationale was that high and low affinity BRET acceptor ligands may enable our sensing window to be shifted into different concentration regimes (e.g. micromolar versus nanomolar), increasing the utility of our method.

**Figure 2.**
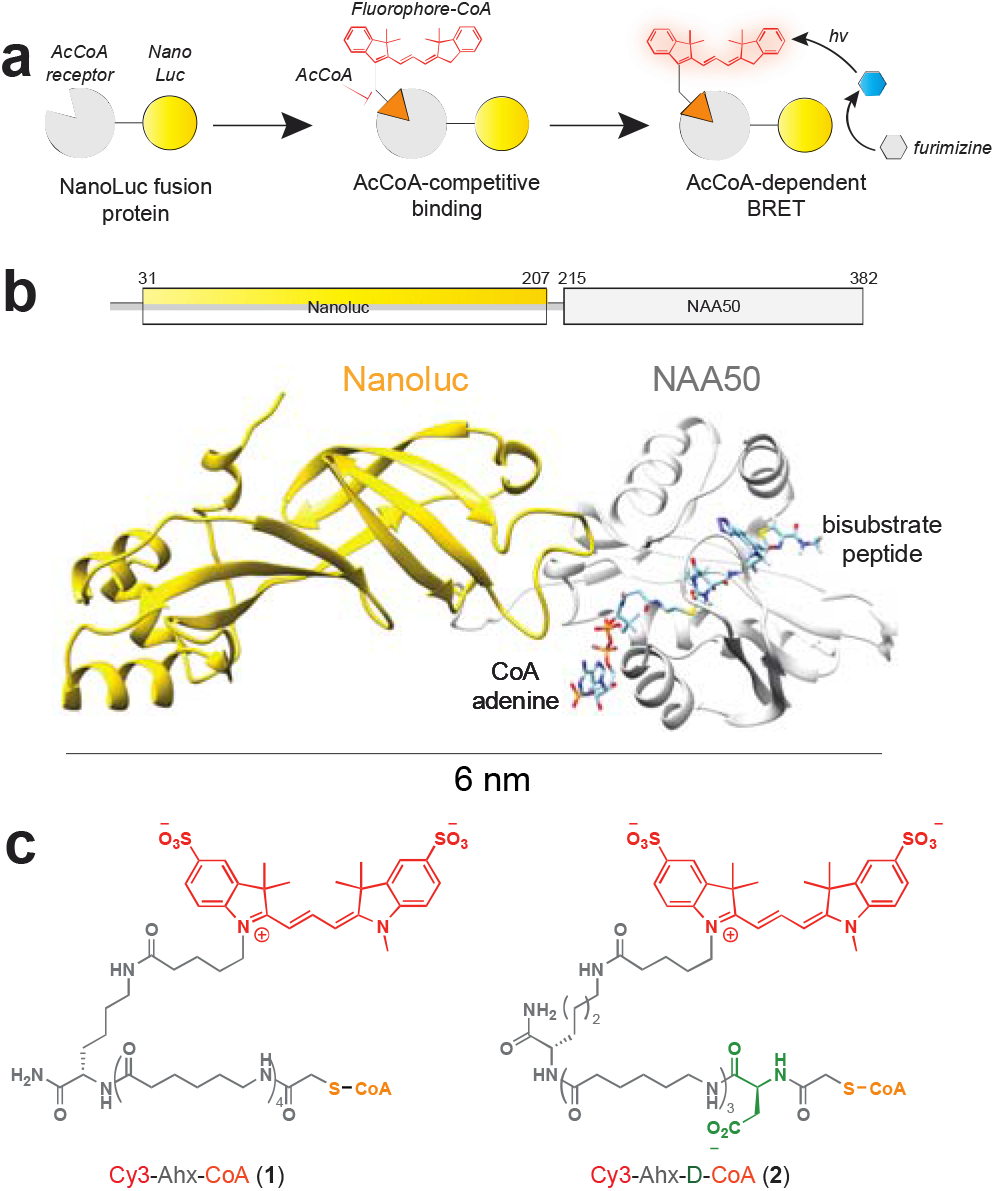
(a) Competitive BRET assay. (b) Domain architecture and predicted structure of Nanoluc-NAA50. CoA bisubstrate is positioned based on a prior NAA50 structure (PDB: 6wf3). (c) Structures of CoA BRET acceptors **1-2**.

To facilitate these studies Nanoluc-NAA50 was recombinantly expressed and purified (Fig. S4). CoA analogues were synthesized via a previously reported method (Scheme S1).^34^ Chemoproteomic capture confirmed preferential binding of acetyl-CoA to the fusion protein (Fig. S5). To assess this system’s capabilities as a BRET reporter (Fig. 3a), purified NanoLuc-NAA50 (Fig. 3b) was incubated with the Nanoluc substrate furimazine in the presence of increasing Cy3-Ahx-CoA (**1**). Evaluation of BRET by comparing the ratio of Cy3 (λ_em_ =590 nm) to furimazine (λ_em_ =460 nm) emission (Fig. 3c) revealed a dose-dependent increase in signal, with a maximum change of ~15-fold observed in the presence of 100 nM **1** (Fig. 3d). BRET signal intensity decreased slightly after 30 minutes (Fig. S6), as expected due to consumption of furimazine substrate. These studies establish the capability of Cy3-CoA **1** to serve as a BRET acceptor for Nanoluc-NAA50.

**Figure 3.**
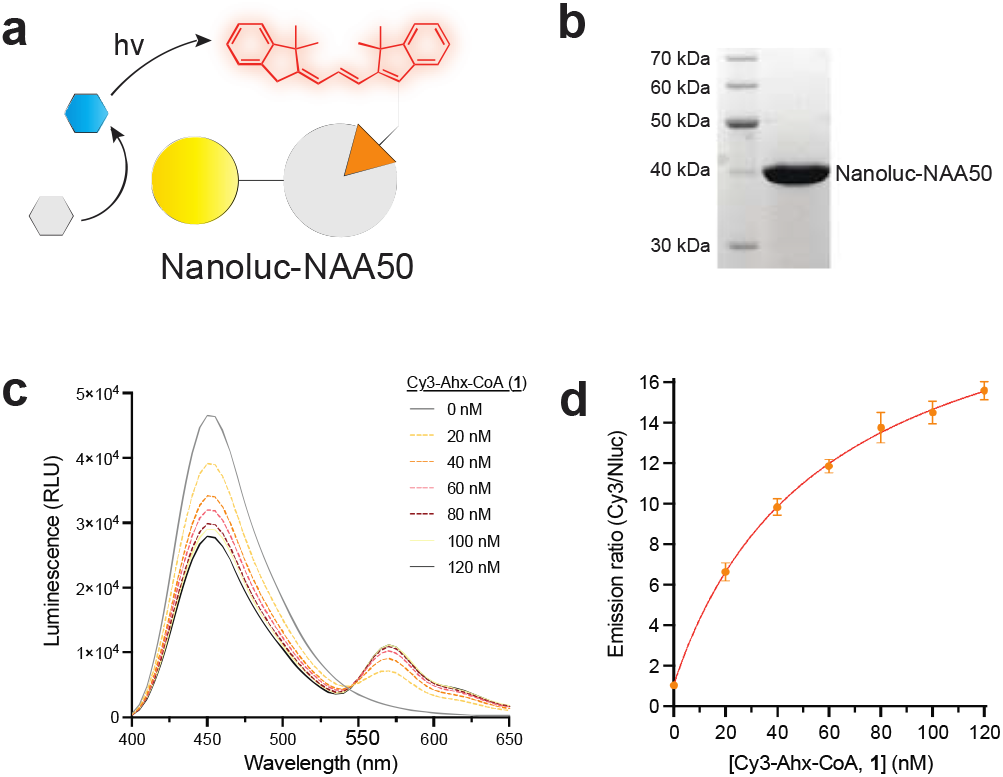
(a) BRET assay schematic. (b) SDS-PAGE of purified Nanoluc-NAA50. (c) Emission spectra of Nanoluc-NAA50 treated with furimazine in the presence of increasing **1**. n=3 replicates. (d) BRET emission ratio of Nanoluc-NAA50 treated with furimazine in the presence of increasing concentrations of **1**. n=3 replicates.

Next, we defined the properties and tunability of this system as a ligand-displacement assay for acetyl-CoA. Nanoluc-NAA50 was pre-incubated with increasing concentrations of CoA metabolites, followed by addition of Cy3-Ahx-CoA (**1**) and furimazine. Consistent with chemoproteomic data, BRET analysis indicated differential competition of NAA50 occupancy by acetyl-CoA (IC_50_ =61 nM) and CoA (IC_50_ =680 nM; Fig. 4a). To extend our BRET sensing method into higher concentration regimes we tested metabolite competition in the presence of **2**, a CoA ligand with a higher affinity for NAA50.^33^ As anticipated, competition of the tighter-binding CoA BRET acceptor requires higher concentrations of acetyl-CoA (IC_50_ = 620 nM) and CoA (IC_50_ = 7300 nM; Fig. 4b). However, the selectivity for these metabolites remains unchanged. The ability to tune concentration window using structural analogues of the BRET acceptor offers a unique alternative to mutational optimization and - given the facile synthesis of high affinity CoA bisubstrates for NAA50 - suggests the possibility of extending this method across the range of acetyl-CoA concentrations that occur in cells.^1,11^ Assessing the selectivity of our assay (using Cy3-Ahx-CoA **1**) across a wider range of CoAs revealed only the short chain fatty acid metabolites acetyl- and propionyl-CoA significantly competed BRET signal at 1 μM (Fig. 4c-d). While the concentration of acetyl-CoA in cells is typically far higher than propionyl-CoA, the latter can accumulate upon amino acid catabolism,^11^ inborn errors of metabolism,^35^ or microbiome-derived production.^36^ These findings establish a ligand-tunable bioluminescent indicator displacement assay for acetyl-CoA and indicate the primary determinant of specificity is the metabolite host.

**Figure 4.**
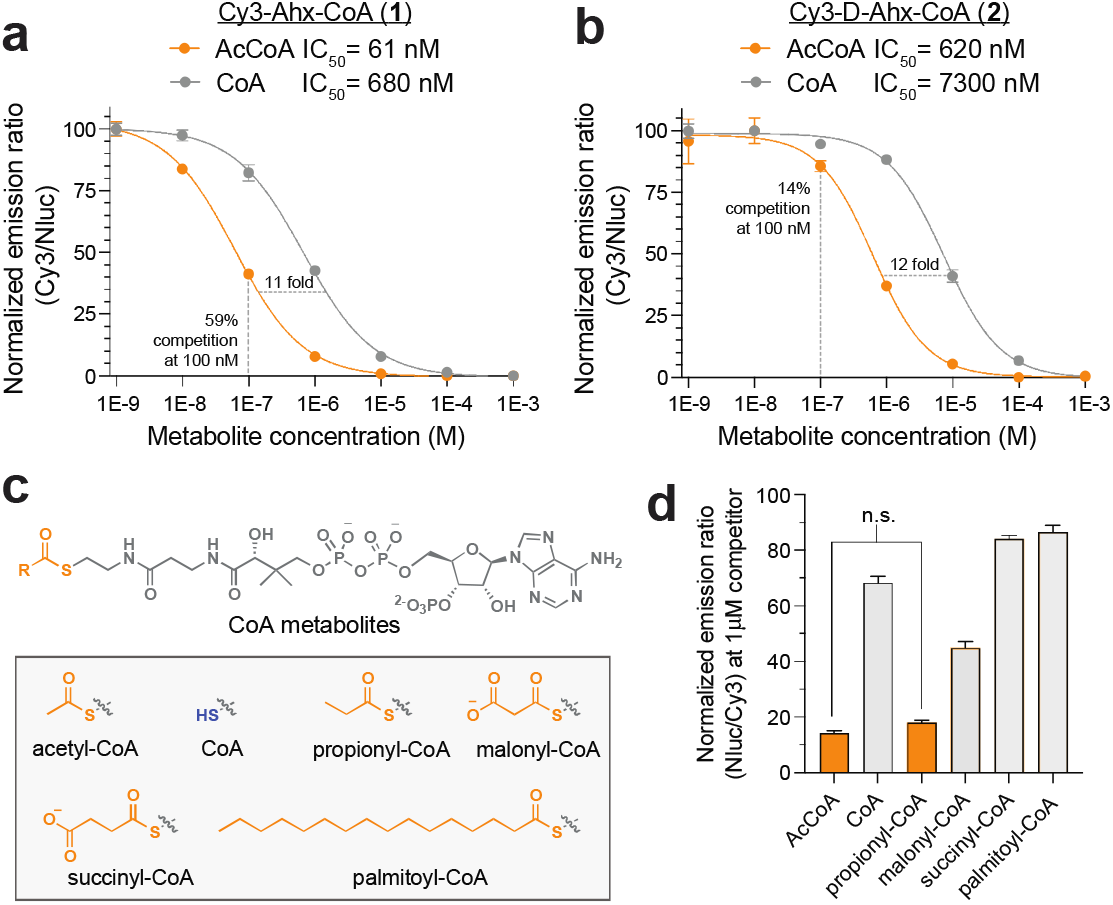
BRET-based detection of CoA metabolites. (a) Competition of **1** by acetyl-CoA (orange) and CoA (gray). n=3. (b) Competition of **2** by acetyl-CoA and CoA. n=3. (c) Structures of CoA metabolites. (d) Competition of Nanoluc-Naa5 0-**1** BRET signal by structurally diverse CoA metabolites at 1 μM. n=3. n.s. = not significantly different (Student’s t-test, twotailed).

Prior work has demonstrated that BRET metabolite sensors can be coupled to a wide range of enzyme assays to facilitate research and clinical diagnostics,^37–39^ a capability that could be extended by a method for detecting acetyl-CoA. To assess whether the modest selectivity window afforded by NAA50 could accommodate such an application, we tested the ability of our BRET ligand-displacement assay to monitor acetyltransferase activity (Fig. 5a). For these assays we incubated the histone acetyltransferase EP300 (75 nM) with acetyl-CoA (1 μM) histone peptide substrate (50 μM), and all components necessary for the BRET detection of acetyl-CoA (Nanoluc-NAA50, furimazine, and **1**). Our expectation was that high concentrations of acetyl-CoA present at the start of the assay would compete with **1** for binding to the Nanoluc-Naa50 donor, inhibiting BRET (Fig. 5a, left). Transfer of acetyl-CoA to the histone peptide by EP300 relieves this competition, enabling **1** to bind and produce a BRET signal (Fig. 5a, right). Pilot studies to establish practicality indicated acetyl-CoA competes Nanoluc-NAA50/**1** BRET in the presence of micromolar CoA (Fig. S7). As anticipated, performing the EP300 enzyme reaction in the presence of ligand-displacement assay components resulted in time-dependent BRET signal production (Fig. 5b). Addition of CPI-1612, a high affinity small molecule inhibitor of EP300,^40^ led to a dose-dependent decrease in BRET signal (IC_50_ = 35 nM, Fig. 5c). Control experiments confirmed CPI-1612 did not interfere with the BRET assay itself (Fig. S8). This method was further extended to monitor the enzymatic activity and small molecule inhibition of KAT6A, another histone acetyltransferase drug target (Fig. S9).^41,42^ While consumption of acetyl-CoA has been monitored as a surrogate for acetyltransferase activity using fluorogenic reactions^43^ or LC-MS,^44^ these assays provide endpoint rather than real-time readouts afforded by the BRET method. Overall, these studies illustrate how selective molecular recognition of acetyl-CoA can be coupled to enzyme activity to develop biochemical assays with novel capabilities.

**Figure 5.**
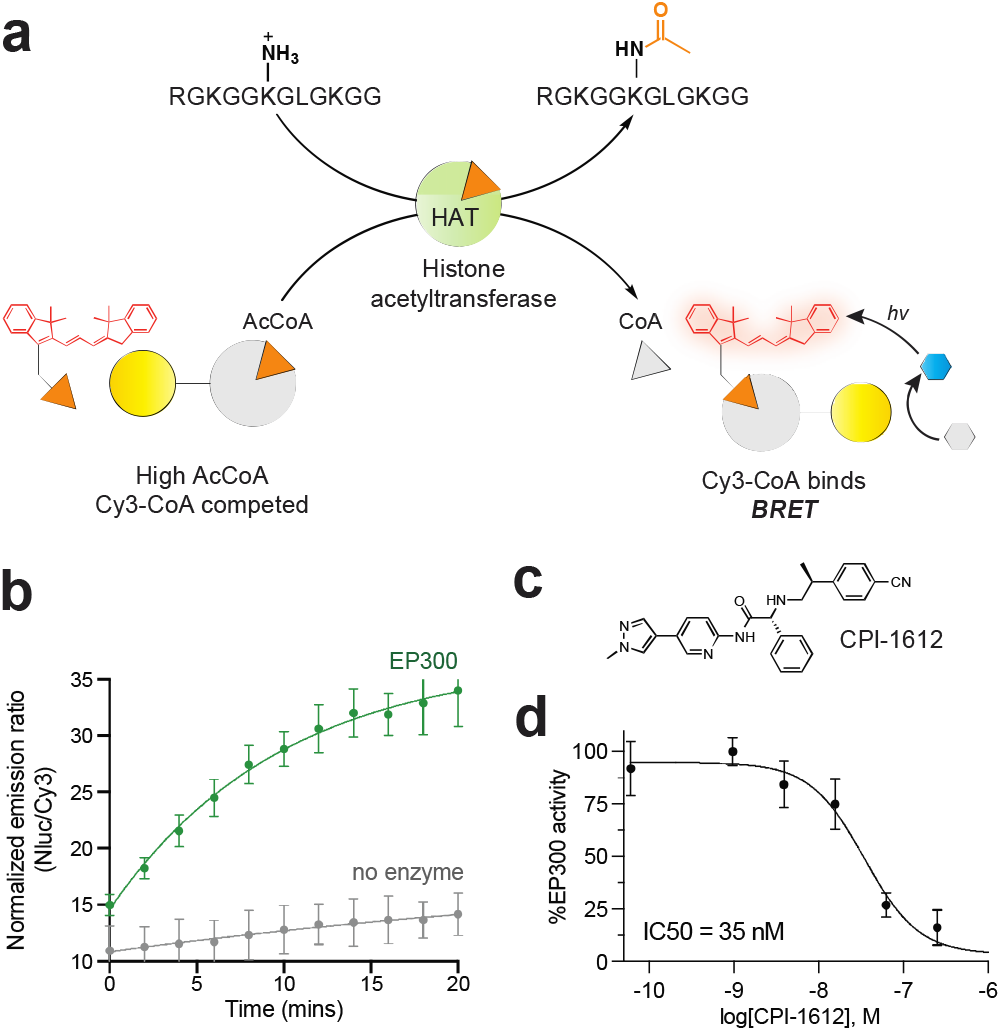
(a) Overall assay schematic. (b) Time-dependent activity of EP300 histone acetyltransferase. (c) Structure of CPI-1612. (d) Dose-dependent inhibition of EP300 by CPI-1612. n=5 replicates.

The selective molecular recognition of acetyl-CoA versus CoA presents a challenge to sensor design due to a lack of scaffolds capable of selective molecular recognition. As a first step to address this challenge, here we describe the chemoproteomics-driven discovery of a selective molecular host for acetyl-CoA. Our approach harnesses the observation that capture of NAA50 by chemoproteomic affinity resins is competed more efficiently by acetyl-CoA than CoA.^10,23^ This enabled the development of a BRET-based indicator displacement assay that can differentiate acetyl-CoA and CoA in a narrow sensing window, whose absolute concentration range can be tuned via structural editing of the Cy3-CoA acceptor ligand, and which can be used for real-time measurements of histone acetyltransferase enzyme activity and small molecule inhibition. Widening the window with which acetyl-CoA can be differentiated from CoA will be important to extend the imaging and diagnostic applications of this approach. Almost all characterized acetyltransferases interact with acetyl-CoA via a binding cleft formed between the β4 and β5 strands that includes a β-bulge in β4 which orients backbone amides towards the carbonyl thioester of acetyl-CoA (Fig. S10).^25^ Prior studies have found mutations in β4, β5, and the adjacent loops can alter acyl-CoA/acetyltransferase interactions,^45–47^ providing a starting point for mutational optimization. Harnessing these optimized receptors for acetyl-CoA imaging will also require further engineering, for example via employment of the Snifit concept (Fig. S11).^16^ Overall, our studies demonstrate how insights from chemoproteomics can address challenges in molecular recognition and provide an analysis workflow and proof-of-concept binding scaffold for target engagement and selective metabolite detection applications.

## Supporting information

Supporting Tables - Excel Format

## ASSOCIATED CONTENT

### Supporting Information

Supporting Information including supplementary figures, tables, and full experimental data available free of charge on the web.

### AUTHOR INFORMATION

**Corresponding Author**

*

## ACKNOWLEDGMENT

The authors thank Dr. Martin Schnermann (NCI) for helpful discussions, Dr. Euna Yoo (NCI) for access to instrumentation, McKenna Crawford (NCI) for proofreading, and Lauren Beaumont (Leidos Inc.) for experimental protocols. We gratefully acknowledge Dr. Kryzsztof Krajewski of the UNC High-Throughput Peptide Synthesis Core facility for peptide synthesis. This work was supported by the Intramural Research Program of the NIH, National Cancer Institute, Center for Cancer Research (ZIA BC011488). In addition, this project has been funded in whole or in part with federal funds from the National Cancer Institute, National Institutes of Health, under contract number HHSN261200800001E.

