## Supplementary Information for "Chemoproteomics yields a selective molecular host for acetyl-CoA"

**Table of Contents for Supporting Information**

**Page**

Supplementary Figures	S2

Supplementary Tables	S13

Supplementary Schemes	S16

Sequence of Nanoluc-NAA50	S17

General materials and synthetic methods	S18

Synthesis of Cy3-CoA BRET acceptors	S19

Expression and purification of Nanoluc-NAA50	S20

Chemoproteomic methods	S21

BRET detection method	S22

BRET acetyltransferase assay	S23

Full gels and Western blot images	S24

References	S25

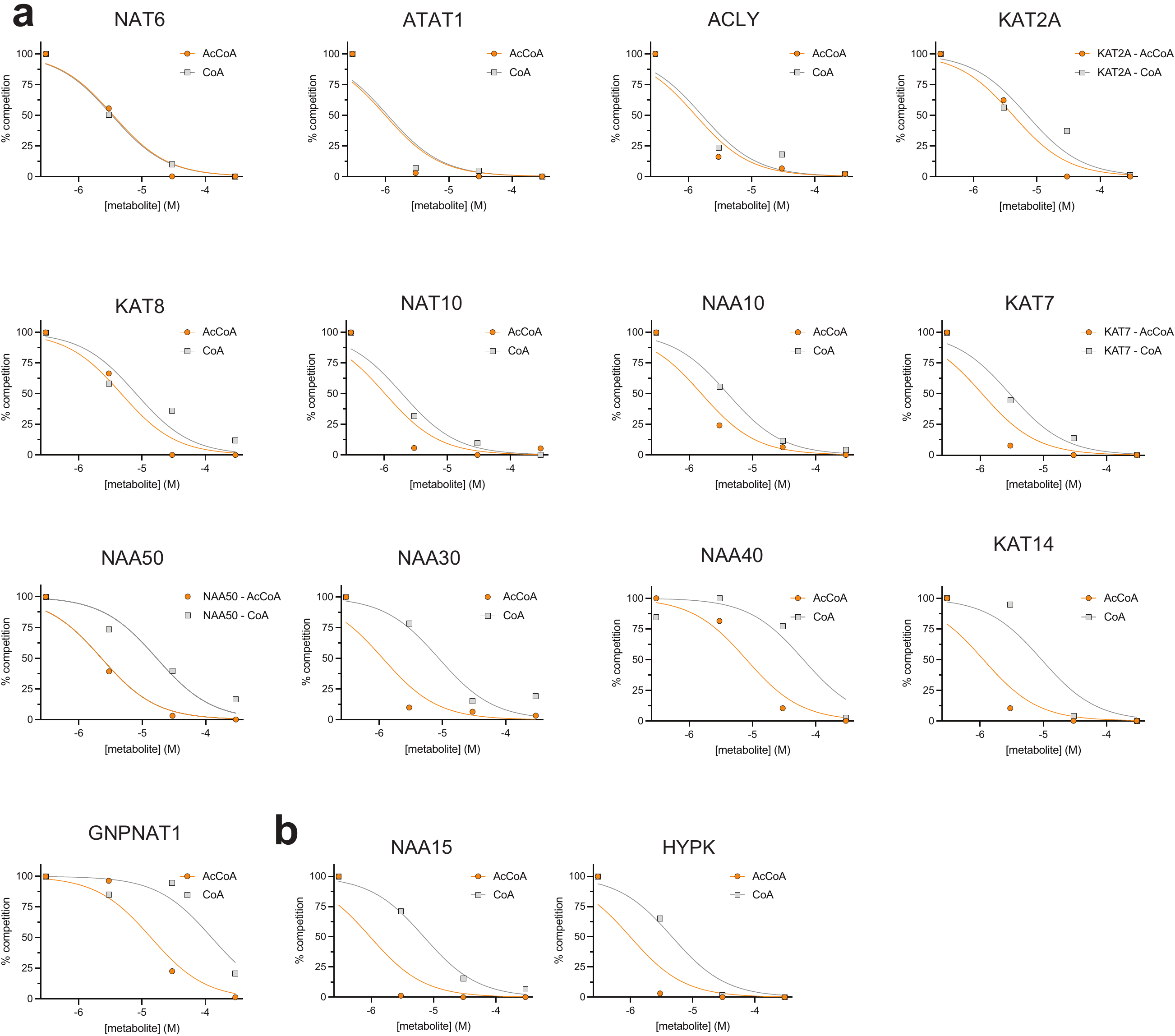

**Figure S1**. (a) Competitive chemoproteomic LC-MS/MS capture data for proteins known to directly bind to acetyl-CoA.^1^ Proteins plotted showed dose-dependent changes (R^2^ ≥ 0.85) in abundance as assessed by the label-free metric distributed normalized spectral abundance factor (dNSAF) after pre-incubation of cell lysates with acetyl-CoA (orange) or CoA (gray) at 0, 3, 30, and 300 μM. Individual points represent averages of triplicate analyses normalized to the no competitor sample (set to 100%) while the sigmoidal represents nonlinear fit of abundance data. (b) Identical analysis for the NAA50 binders NAA15 and HYPK. Numerical values are provided in Excel format as Supporting Information. Unfiltered proteomic data is available via the reference study^1^ and PRIDE (dataset identifier PXD013157).

**Figure S2**. (a) Domain architecture of Gcn5-N-acetyltransferase domain protein NAA50. Arrows represent beta sheets, cylinders represent alpha helices. (b) Structure of NAA50 (PDB: 6wfo) specifying elements involved in acetyl-CoA binding. (c) Structure of NAA40 (PDB: 4u9v) specifying elements involved in acetyl-CoA binding.

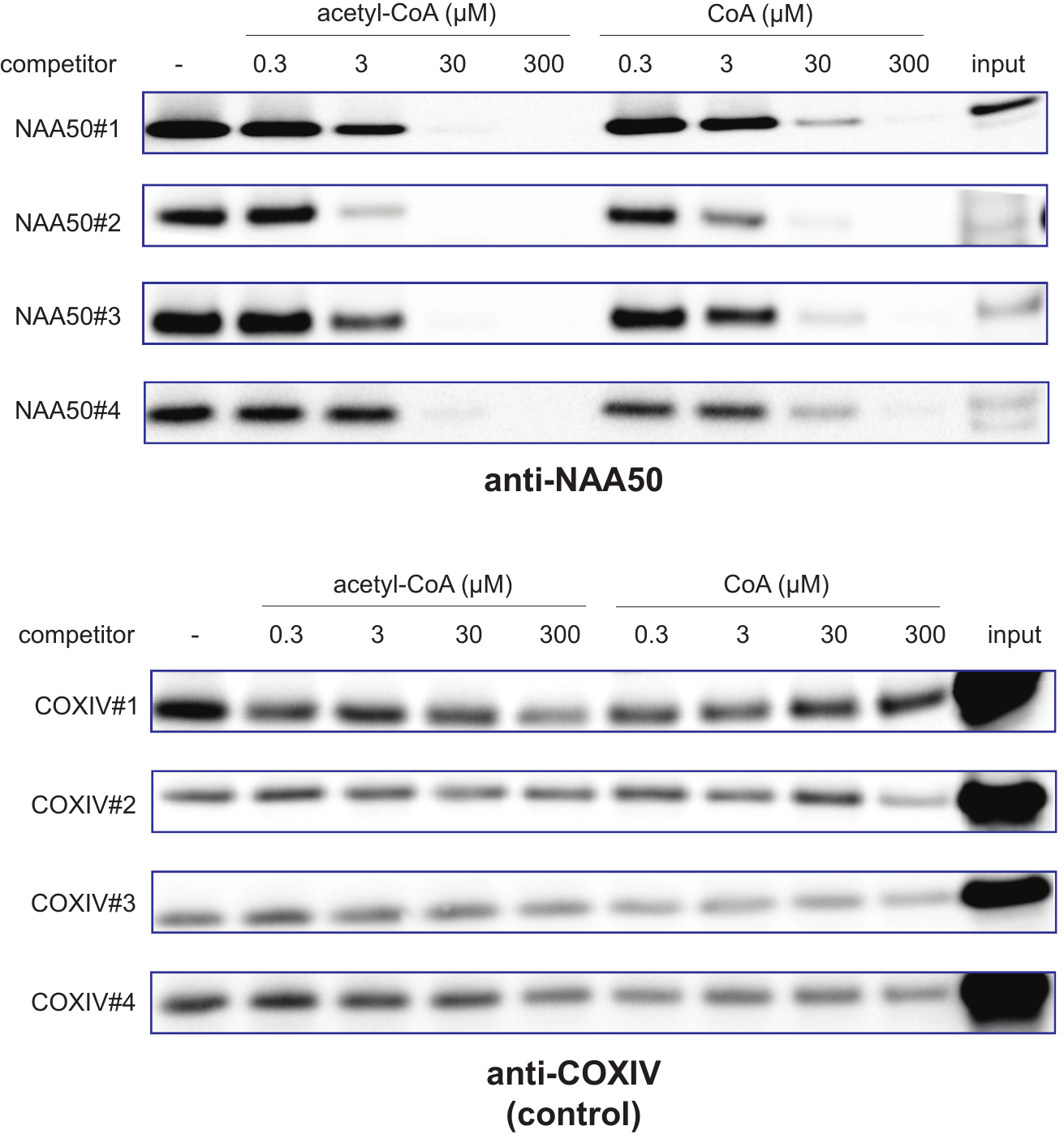

**Figure S3**. Chemoproteomic capture of NAA50 (top) by Lys-CoA resins from HeLa cell lysates is competed by acetyl-CoA and CoA. Increased competition by acetyl-CoA relative to CoA is readily apparent at 30 μM. The non-specific capture target COXIV (bottom) is not competed by the two metabolites. Reprinted with permission from supplementary information, *J. Am. Chem. Soc.* 2016, 138, 20, 6388–6391.^2^

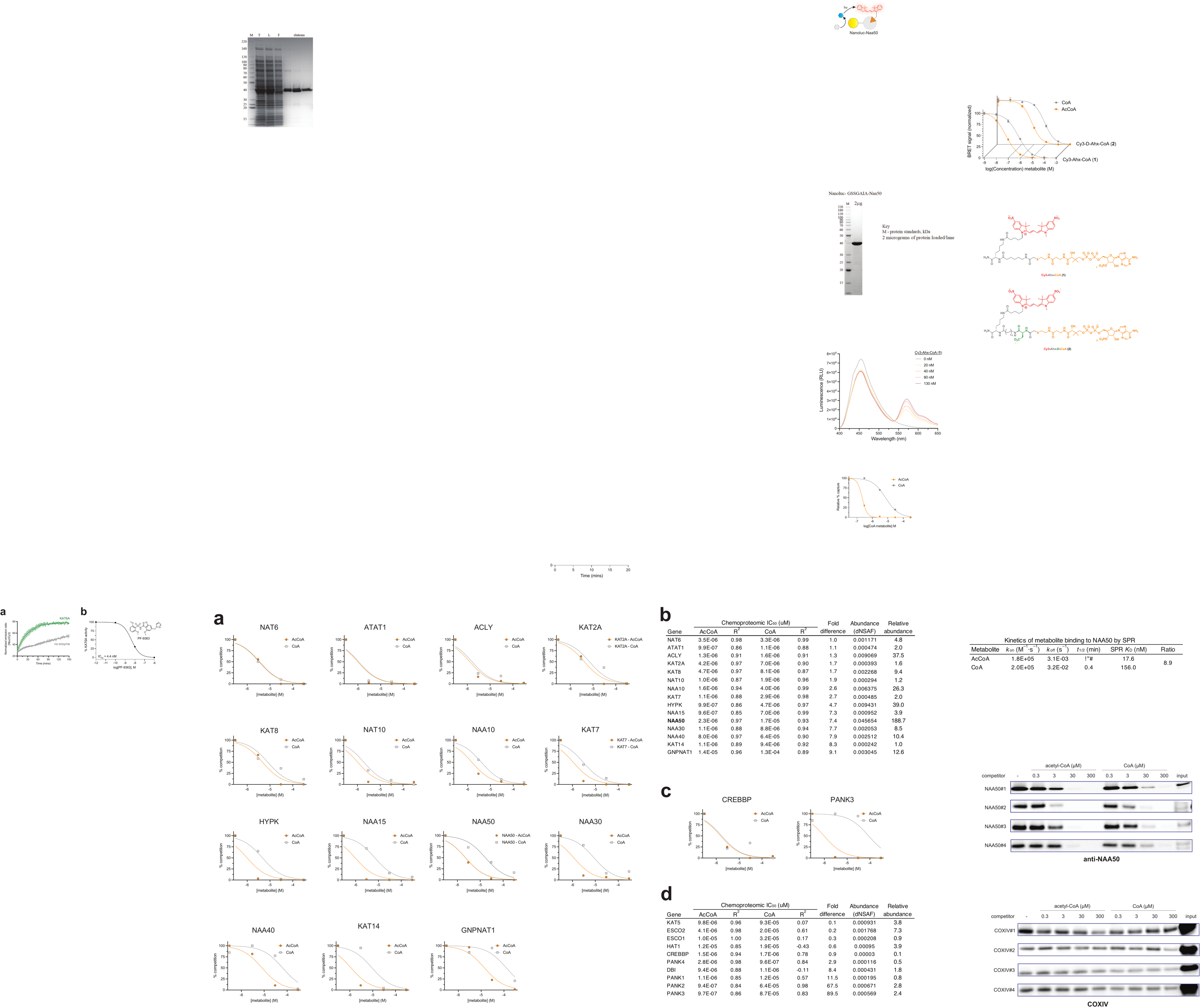

**Figure S4**. Overexpression and purification of Nanoluc-NAA50 using immobilized metal affinity chromatography. M = protein standards, kDa. T = total cell lysate. L = column load (soluble protein). F = column flow through. Elutions = IMAC column elutions (increasing imidazole).

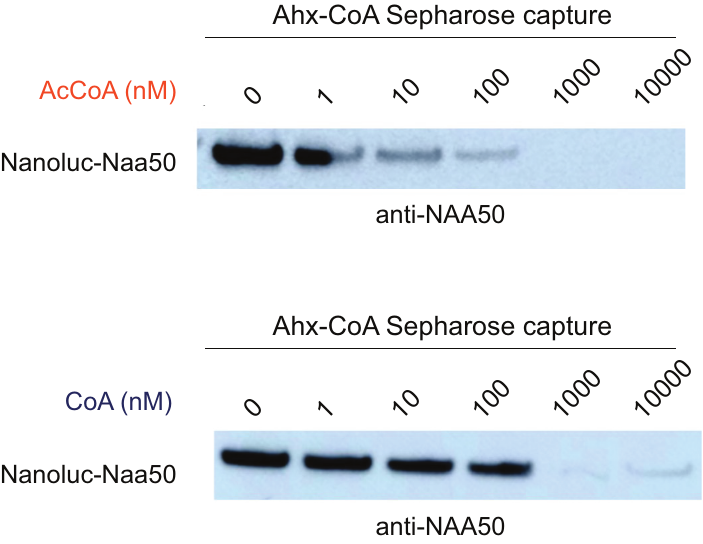

**Figure S5**. Chemoproteomic capture of recombinant Nanoluc-NAA50 by Ahx-CoA resins is competed by acetyl-CoA (top) and CoA (bottom).

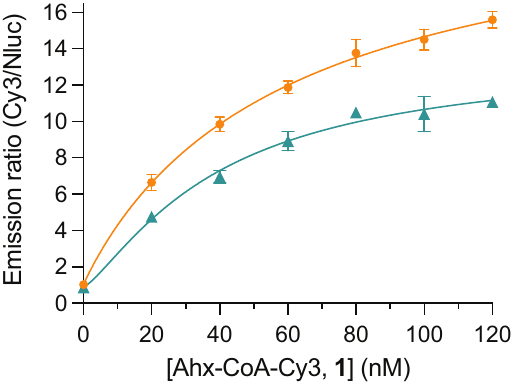

**Figure S6**. BRET emission ratio of Nanoluc-NAA50 treated with furimazine in the presence of increasing concentrations of **1**. Orange = initial values after addition of furimazine (t=0 min). Teal = values 30 min after addition of furimazine (t=30 min). n=3 replicates.

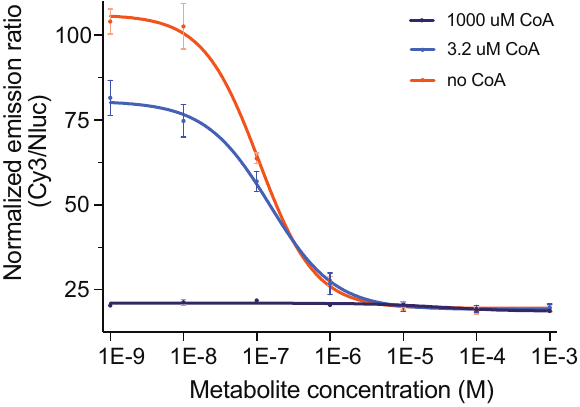

**Figure S7**. Effects of CoA on acetyl-CoA’s ability to competitively inhibit BRET signal production by the Nanoluc-NAA50/**1** complex. Competition is readily observed at 0 and 3.2 μM CoA; however, high concentrations (1000 μM) of CoA interfere with detection.

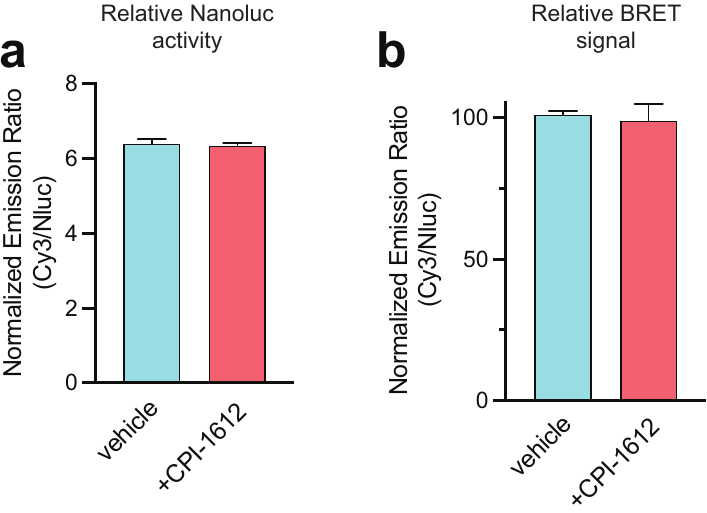

**Figure S8**. (a) Emission data from reaction mixture in which Nanoluc-NAA50 and furimazine were treated with or without CPI-1612 (100 μM). The lack of change in the emission suggests CPI-1612 is not disrupting Nanoluc turnover of furimazine. (b) Emission data from reaction mixture in which Nanoluc-NAA50/**1** and furimazine were treated with or without CPI-1612 (100 μM). The lack of change in the BRET signal suggests CPI-1612 is not disrupting the binding of **1** to Nanoluc-NAA50.

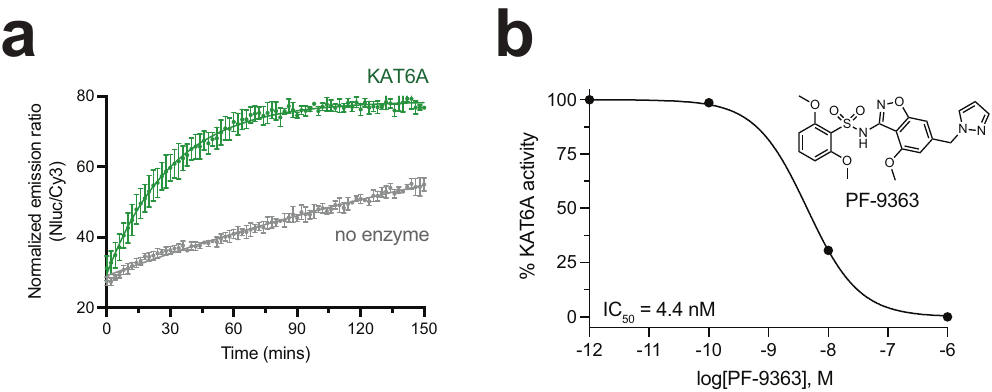

**Figure S9**. (a) Time-dependent activity of KAT6A histone acetyltransferase. (b) Dose-dependent inhibition of KAT6A by PF-9363. n=3 replicates.

**Figure S10**. (a) Domain architecture of Gcn5-N-acetyltransferase domain protein NAA50. Arrows represent beta sheets, cylinders represent alpha helices. (b) Structure of NAA50 (PDB: 6wfo). (c) Structure of NAA40 (PDB: 4u9v). (d) Structure of GNPNAT1 (PDB: 4u9v). (e) Structure of KAT2A (PDB: 4u9v). (f) Structure of EP300 (PDB: 4u9v). Notations specify structural elements (β4 β bulge/β5 Leu) involved in acetyl-CoA recognition. (g) Alignment of structural elements (β4 β bulge/β5 Leu) across multiple acetyltransferases. Residues in blue are directed towards the acetyl-CoA binding site. Mutation of residues shaded in yellow have been shown to affect acetyl-CoA or acyl-CoA binding.

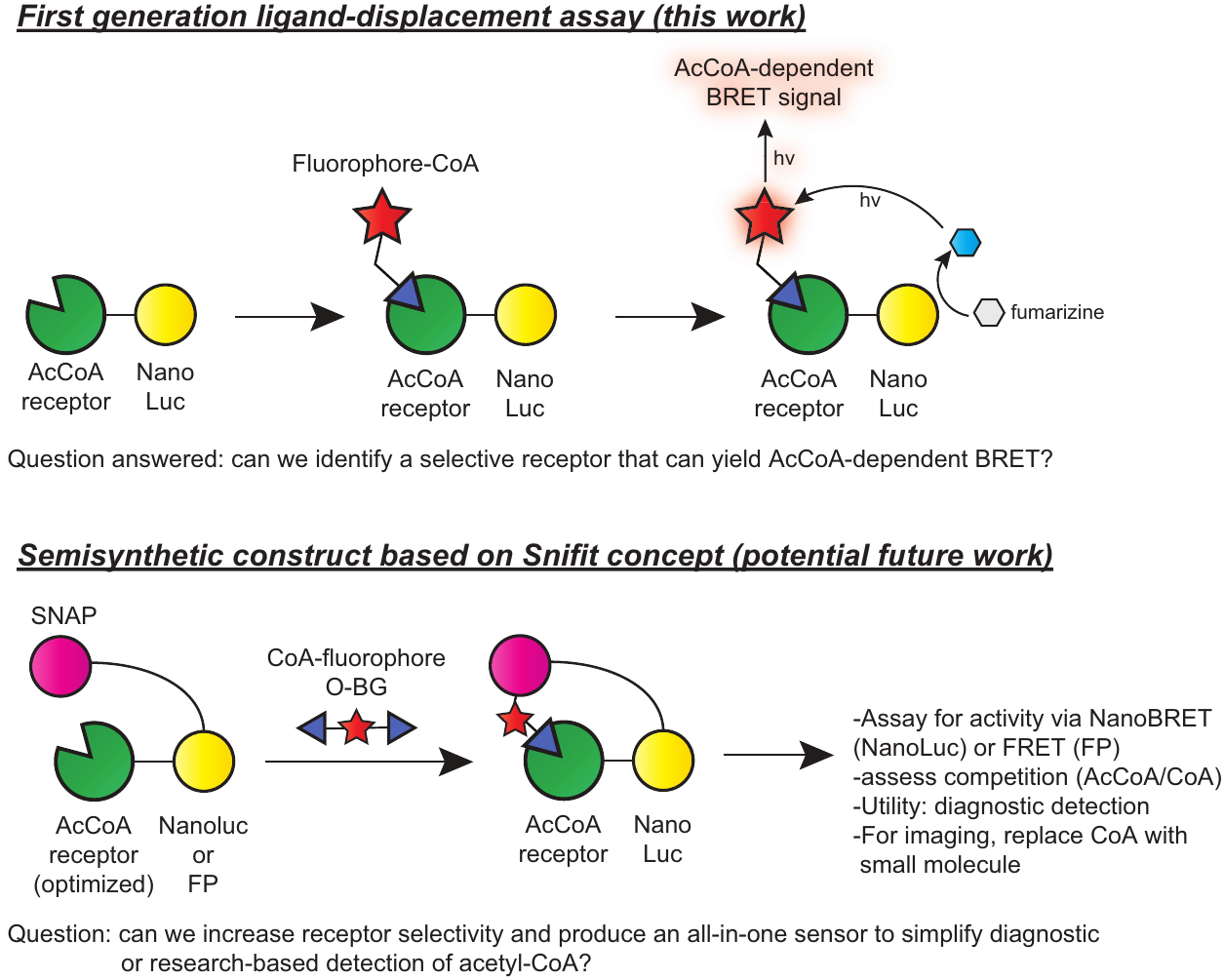

**Figure S11**. Overview of first-generation BRET-based ligand-displacement assay for acetyl-CoA (this work) and potential design elements necessary for employing the fluorescent CoAs and NAA50 host identified here to Snifit (**SN**AP-tag based **i**ndicator with a **f**luorescent **i**ntramolecular **t**ether)-based detection of acetyl-CoA in potential future work.

**Supplementary Tables**

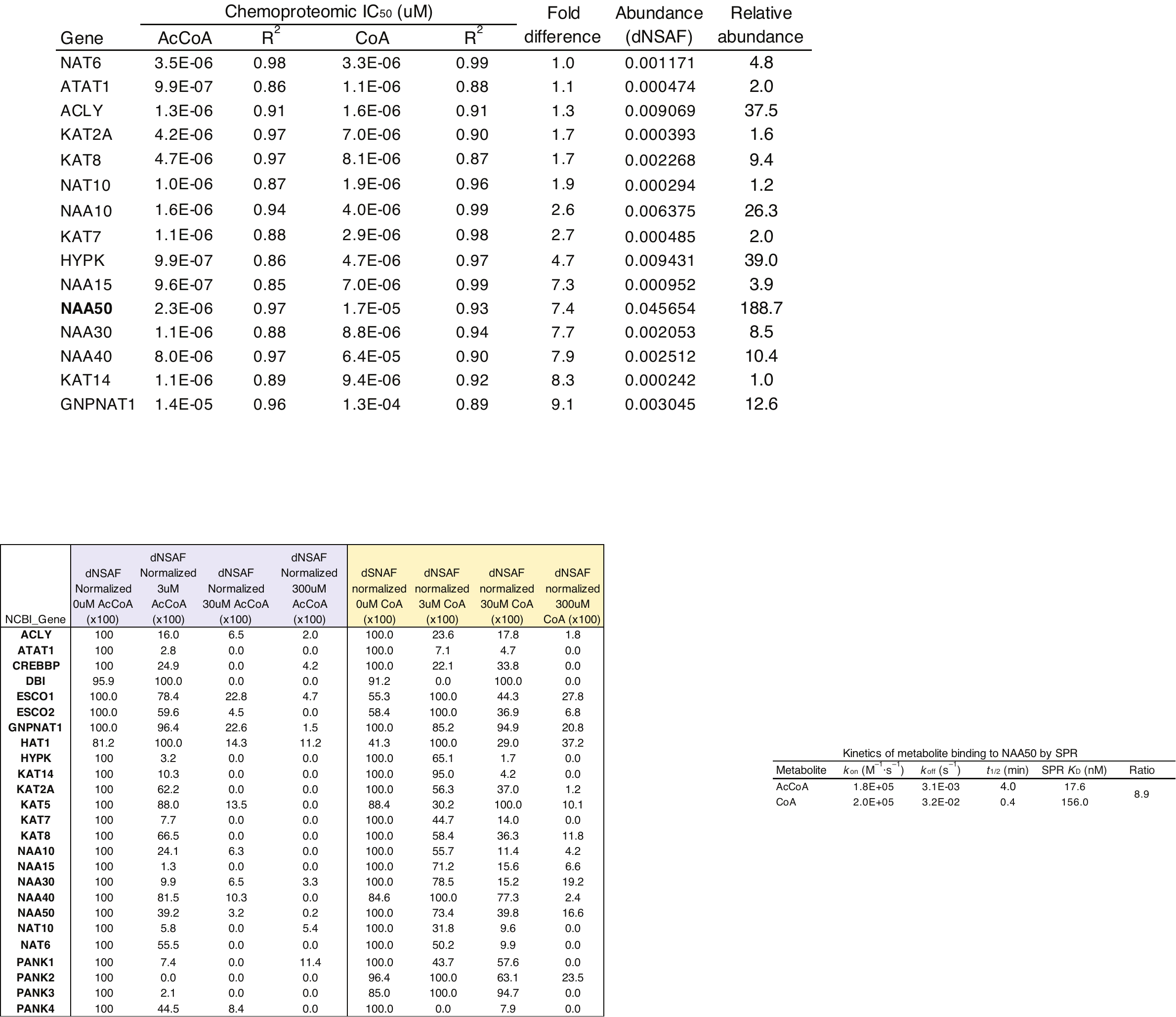

**Table S1.** Competitive chemoproteomic capture of 23 acetyl-CoA-binding proteins from cell lysates by CoA affinity resin.^1^ Protein abundance was quantified by LC-MS/MS using label-free analysis (distributed normalized spectral abundance factor [dNSAF]) after pre-incubation of lysates with acetyl-CoA or CoA (gray) at 0, 3, 30, and 300 μM. Values represent averages of triplicate analyses normalized to the no competitor sample (set to 100%). Numerical values for >1000 proteins captured by CoA affinity resin are provided in Excel format as Supporting Information. Additional proteomic data is available via the reference study^1^ and the PRIDE database (dataset identifier PXD013157). Data for the NAA50 interactors NAA15 and HYPK is also included.

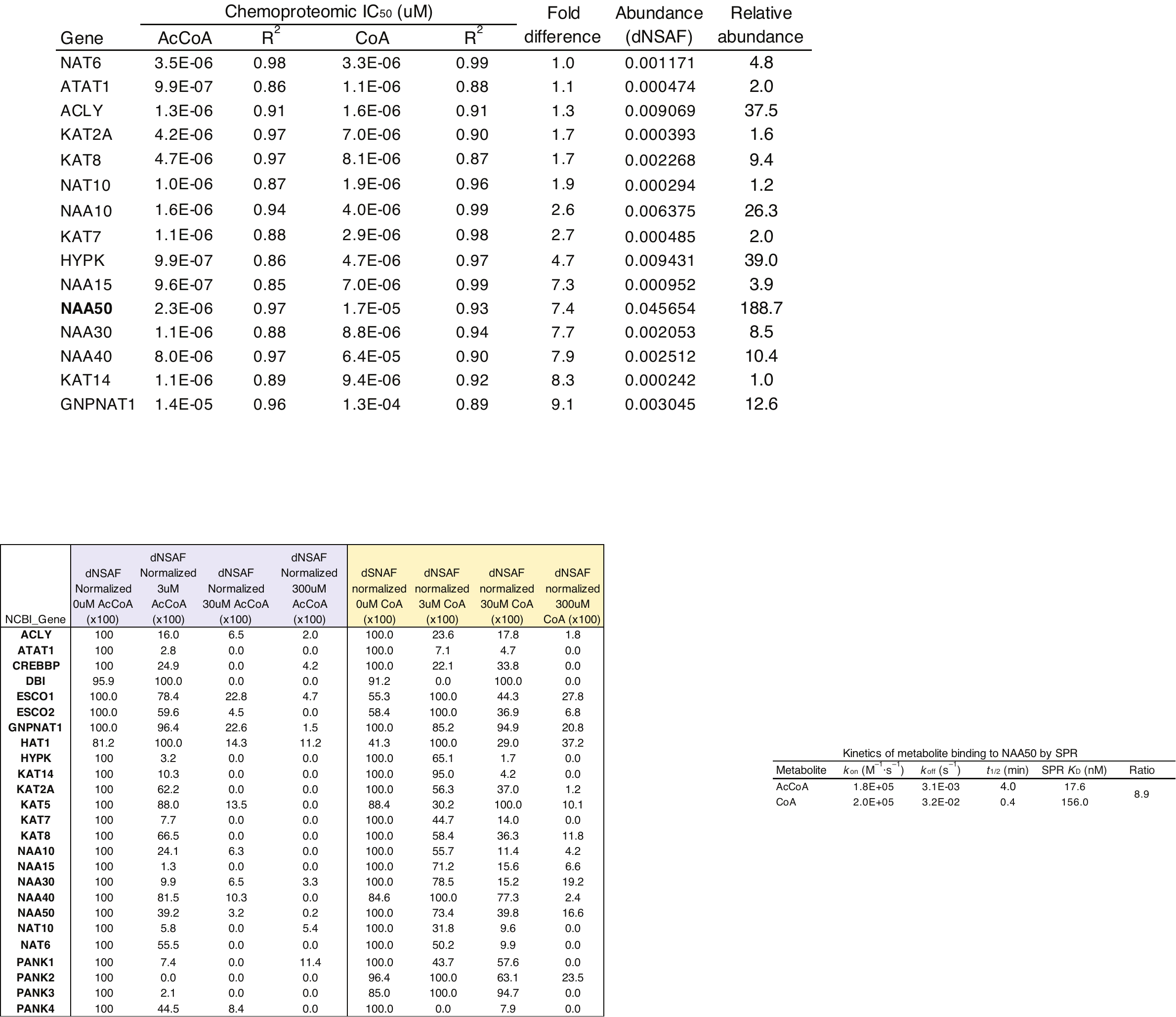

**Table S2.** Normalized dNSAF abundance data (n=3) for proteins captured from HeLa cell lysates by Lys-CoA Sepharose in the presence or absence of the specified metabolites (0, 3, 30, and 300 μM) were fit to a nonlinear regression model using GraphPad Prism to produce IC_50_ and goodness of fit (R^2^) values. ‘Fold difference’ represents the ratio of CoA to AcCoA IC_50_ for each protein. ‘Relative abundance’ represents the average dNSAF for each protein normalized to a relatively weakly captured acetyl-CoA binding protein (KAT14). NAA50 is highlighted in bold.

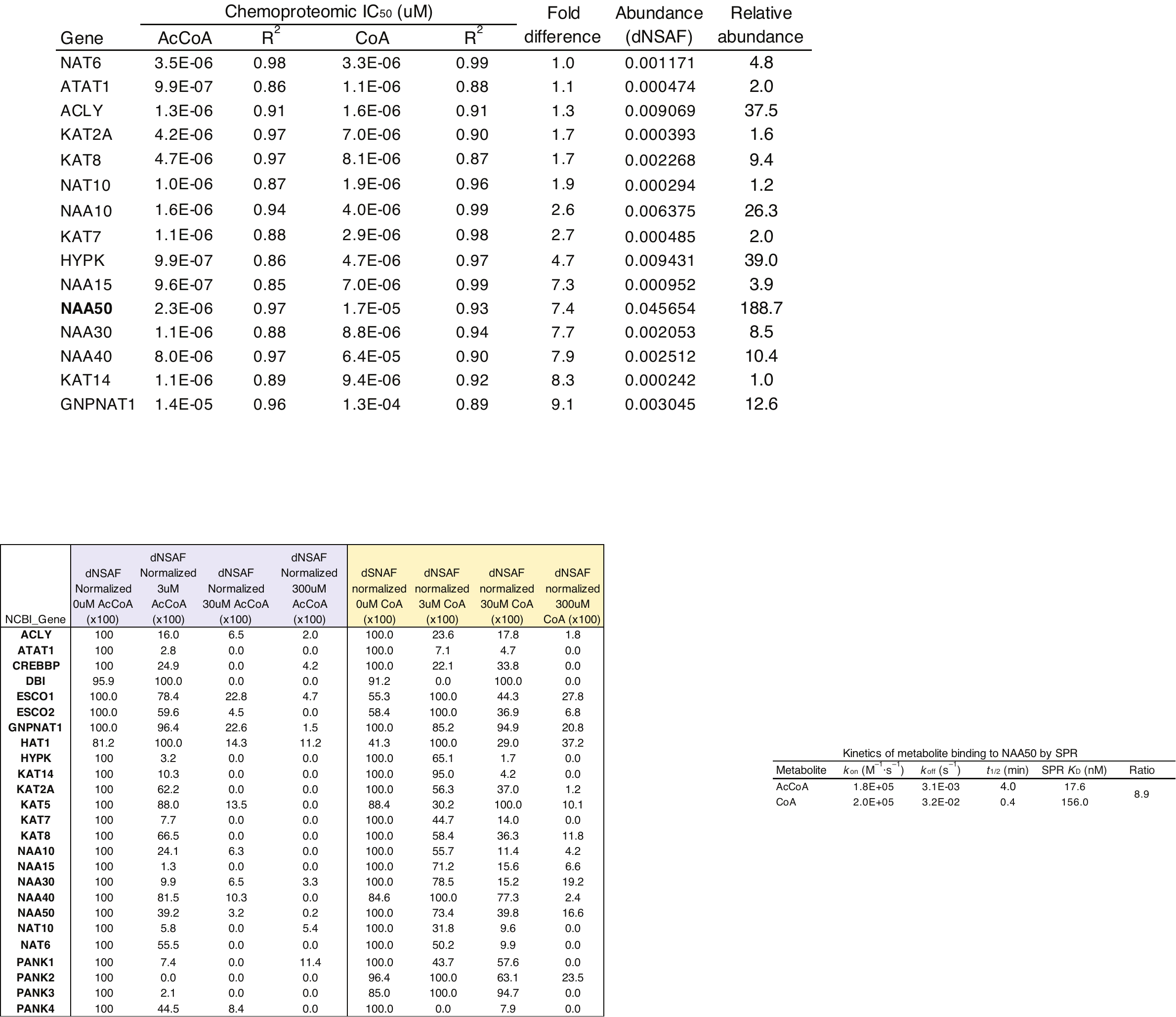

**Table S3.** Surface plasmon resonance (SPR)-based binding kinetics of acetyl-CoA and CoA to recombinant NAA50. Ratio indicates the relative ratios of the Kd values. Reprinted with permission from supplementary information, *ACS Med. Chem. Lett.* 2020*,* 11, 6, 1175–1184.^3^

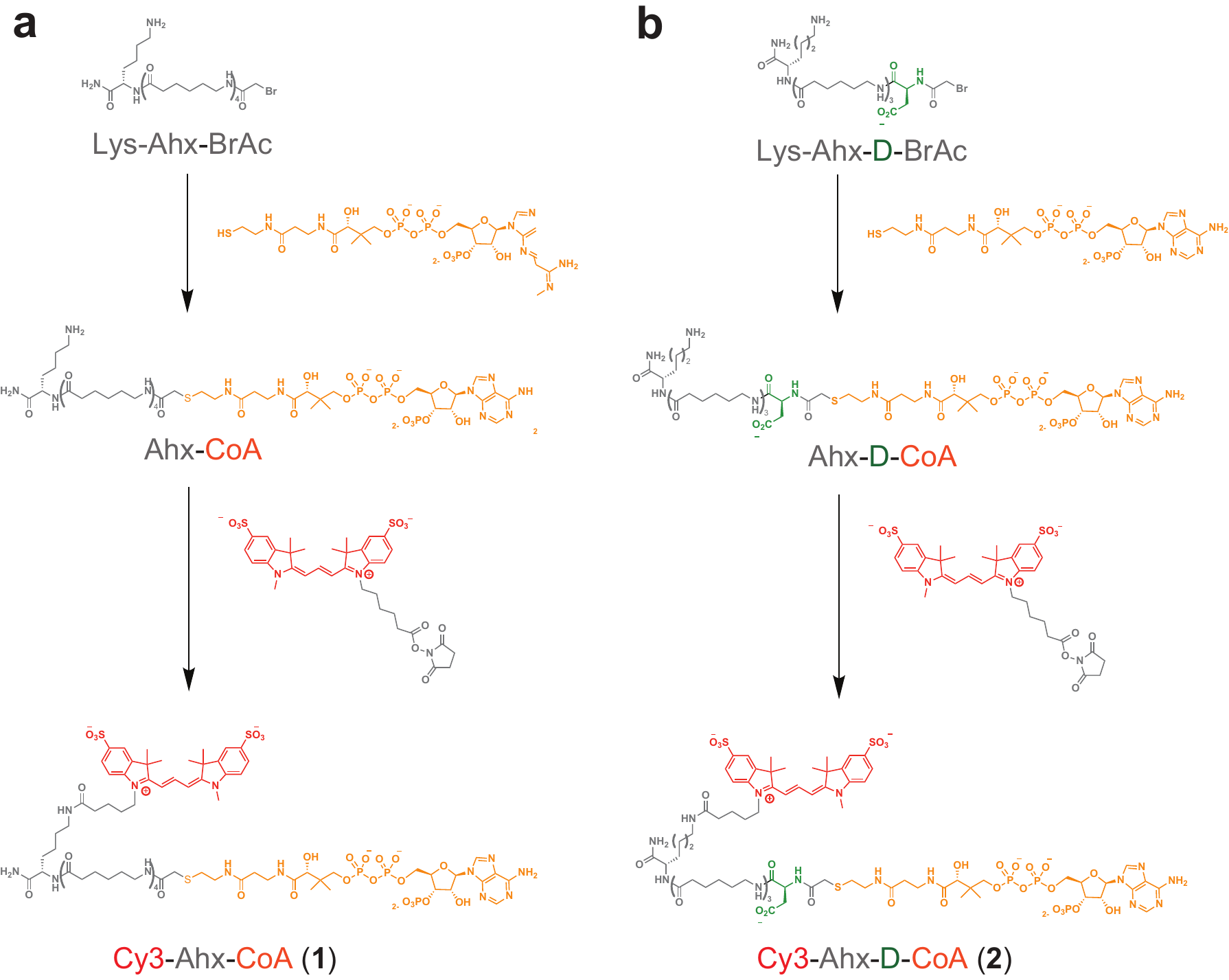

**Scheme S1.** (a) Synthesis of Ahx-Cy3-CoA **1**. (b) Synthesis of D-Ahx-CoA **2**. Bromoacetamide-Ahx_4_ and Bromoacetamide-D-Ahx_3_ precursors synthesized via Rink amide synthesis as previously described.^4^

**Sequence of Nanoluc-NAA50**

MRSGSHHHHHHRSDITSLYKKVGENLYFQGVFTLEDFVGDWRQTAGYNLDQVLEQGGVSSLFQNLGVSVTPIQRIVLSGENGLKIDIHVIIPYEGLSGDQMGQIEKIFKVVYPVDDHHFKVILHYGTLVIDGVTPNMIDYFGRPYEGIAVFDGKKITVTGTLWNGNKIIDERLINPDGSLLFRVTINGVTGWRLCERILAGSGSGSGAIAKGSRIELGDVTPHNIKQLKRLNQVIFPVSYNDKFYKDVLEVGELAKLAYFNDIAVGAVCCRVDHSQNQKRLYIMTLGCLAPYRRLGIGTKMLNHVLNICEKDGTFDNIYLHVQISNESAIDFYRKFGFEIIETKKNYYKRIEPADAHVLQKNLKVPSGQNADVQKTDN

Cyan = TEV-cleavable 6xHis tag

Yellow = Nanoluc

Green = Gly/Ser linker

Grey = NAA50

**General materials and methods**

Unless otherwise specified, all chemicals and solvents were purchased from Sigma, VWR, or Fisher and used without further purification. HPLC-purified Lys-Ahx_4_-bromoacetamide and Lys-Ahx_3_-D-bromoacetamide precursors were purchased from the University of North Carolina High-Throughput Peptide Synthesis Core Facility. Furimazine was purchased from Aobious (AOB36539). Analytical LC-MS of CoA analogues were carried out using an Agilent 1200 Quaternary LC-MS with an Agilent EC-C18 column (2.7 μM, 2.1x50 mm) employing a gradient of 0 → 30% acetonitrile/0.1% formic acid over 6 minutes at a flow rate of 0.6 mL/min. Purification of Cy3-CoAs was performed on an Agilent 1260 Infinity Quaternary HPLC using a Phenomenex Luna C18 column (5 uM, 100 Å) using initial gradient of 0 → 25% acetonitrile over 75 minutes with an increasing flow rate of 2 mL/min → 5mL/min over the same duration was followed by a gradient of 25% → 45% acetonitrile over 20 minutes at a constant flow rate of 5 mL/min. Propionyl-CoA	(#317-66-8) and succinyl-CoA (#108347) were obtained from CoALA Biosciences. Butyryl-CoA (#102282-28-0) and palmitoyl-CoA (#188174-64-3) were obtained from Sigma. Malonyl-CoA (#116928-84-8) was obtained from Santa Cruz Biotechnology. Recombinant EP300 was purchased from Enzo (BML-SE451-010). Recombinant KAT6A was purchased from SignalChem (#K315-381BG). Histone H4 1-21 peptide was purchased from Anaspec (AS-62499), CPI-1612 (#HY-136285) and PF-9363 (#HY-132283) were purchased from MedChemExpress. SDS-PAGE was performed using Bis-Tris NuPAGE gels (4–12%, Invitrogen #NP0321), and MES running buffer (Life technologies #NP0002) in Xcell SureLock MiniCells (Invitrogen) according to the manufacturer’s instructions. For western blotting, SDS-PAGE gels were transferred to nitrocellulose membranes (Novex, Life Technologies # LC2001) by electroblotting at 30 volts for 1 hour using a XCell II Blot Module (Novex). Membranes were blocked using StartingBlock (PBS) Blocking Buffer (Thermo Scientific) for 20 minutes, then incubated overnight at 4 °C in a solution containing the primary antibody to NAA50 (16120-1-AP, Protein Tech, 1:1000 dilution) in the above blocking buffer with 0.05% Tween 20. The membranes were next washed with TBST buffer and incubated with a secondary HRP-conjugated antibody (anti-rabbit IgG, HRP-linked [7074], Cell Signaling, 1:1000 dilution) for 1 hour at room temperature. The membranes were again washed with TBST, treated with chemiluminescence reagents (Western Blot Detection System, Cell Signaling) for 1 minute, and imaged for chemiluminescent signal using an ImageQuant 800 Digitial Imaging System (Cytiva). Total protein content on western blots was visualized by Ponceau stain. Optical measurements were recorded on a Cytation 5 Multimode Plate Reader (Biotek).

**Synthesis of Cy3-CoA BRET acceptors**

Lys-Ahx_4_-bromoacetamide (0.013 mmol) was dissolved in 100 mM pH 8 NaHCO_3_ (0.5 mL) and to this solution was added Coenzyme A (10 mg, 0.013 mmol). The solution was stirred for 30 minutes, diluted with aqueous 0.1% TFA (5 mL), and the pH adjusted to 2 using dropwise addition of 1M HCl. The solution was directly purified via preparative HPLC and lyophilized to provide the product which we refer to as Ahx-CoA (Scheme S1) as a fluffy white solid. The CoA analogue was quantified by the method of Killenberg and Dukes using the molar extinction coefficient (ε) for CoA of 15,000 M^-1^ cm^-1^ at λmax of 259 nm. This material was used for the preparation of chemoproteomic capture resins (vide infra) or carried forward for fluorophore conjugation. To synthesize Cy3-Ahx-CoA (**1)**, Ahx-CoA (5 mg, 0.035 mmol) was dissolved in anhydrous DMF and three equivalents of *N,N*-diisopropylethylamine (18.4 μL, 0.105 mmol) was added. Separately, Sulfo-Cyanine3 NHS Ester (Lumiprobe #21320) was dissolved in 0.1 mL DMSO. The two solutions were combined, vortexed for 10 seconds, and incubated at room temperature overnight protected from light. Upon completion of the reaction as assessed by analytical LC-MS, the solution was diluted in 0.1% TFA/H_2_O (5 mL) and acidified to pH 2 using 1M HCl. Cy3-Ahx-CoA (**1**) was purified by preparative HPLC and lyophilized to provide the product as a red solid. Cy3-Ahx-CoA (**2**) was prepared by an identical procedure starting from Lys-Ahx_3_-D-bromoacetamide precursor. The identity and purity of **1** and **2** was verified by analytical LC-MS analysis prior to use in BRET assays.

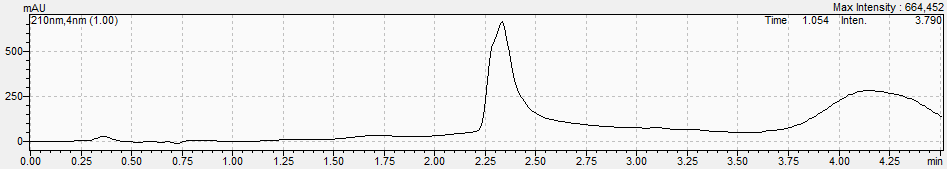

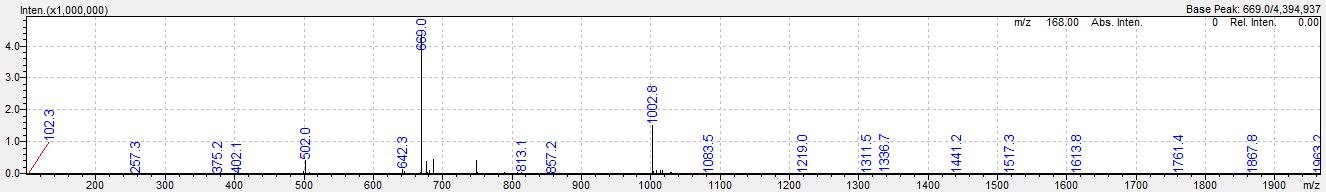

LC-MS trace for Cy3-Ahx-CoA (**1**). [M+2H] C_84_H_134_N_16_O_28_P_3_S_3_ predicted 1001.9, found 1002.8.

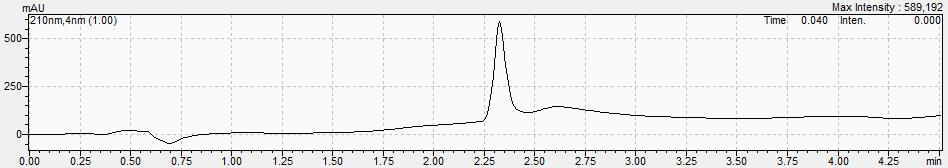

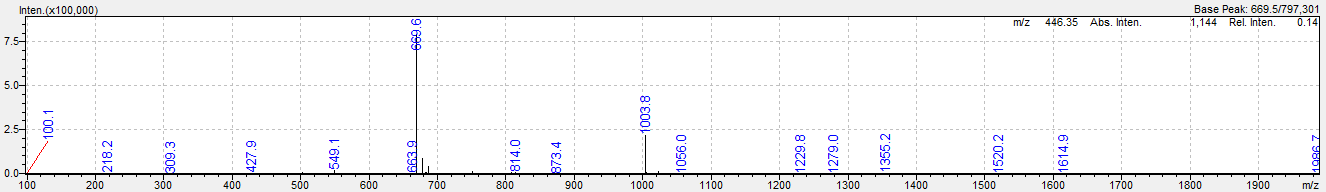

LC-MS trace for Cy3-D-Ahx-CoA (**2**). [M+2H] C_81_H_125_N_16_O_31_P_3_S_3_ predicted 1003.4, found 1003.7.

**Expression and purification of Nanoluc-NAA50**

A Nanoluc-NAA50 expression construct incorporating a TEV-cleavable N-terminal 6xHis-tag was codon-optimized for *E. coli* expression and synthesized by ATUM. Protein was expressed from using the Dynamite expression protocol.^6^ Briefly, a culture of *E. coli* harboring Nanoluc-NAA50 plasmid were grown in non-inducing MDAG-135 medium, overnight at 37 °C. This culture was used to inoculate 2 L of Dynamite media. Cells were grown at 37 °C until an OD600 nm of 6–8 was reached, before induction with 0.5 mM IPTG and incubation at 16 °C for 18–20 hours. Cells were collected by centrifugation, lysed using a microfluidizer, and clarified by ultracentrifugation (100,000xg, 30 minutes at 4 °C). Protein was isolated by immobilized metal affinity chromatography (IMAC) using a Ni Sepharose High Performance column (GE Healthcare) on an NGC chromatography system (Bio-Rad).

**Chemoproteomic methods**

Chemoproteomic LC-MS/MS data analyzed in Figure 1/S1 were derived from a previous dataset^1^ in which HeLa cell proteomes were enriched using Lys-CoA Sepharose in the presence or absence of acetyl-CoA or CoA competitors. Abundance was quantified using distributed normalized spectral abundance factor (dNSAF), a label-free metric that normalizes spectral counts relative to overall protein length with average of the no competition experiments representing 100%. Plotted values represent averages of triplicate analyses normalized to the highest abundance sample (in most cases ‘no competitor,’ set to 100%). Sigmoidal represents fit of abundance data to a nonlinear regression analysis (GraphPad Prism). Numerical values are provided in Excel format as Supporting Information. Additional proteomic data is available via the reference study^1^ and PRIDE (dataset identifier PXD013157).

Ahx-CoA Sepharose resin was prepared using NHS-Activated Sepharose 4 Fast Flow resin (Cytiva, #17-0906-01) following the previously reported method and manufacturer’s protocol.^2,4^ Briefly, resin was washed with ice cold 1 mM HCl (10 volumes), pelleted, and supernatant removed. Next, a 3.4 mM solution of Ahx-CoA (free amine functionalized) was prepared in 1x PBS and added to the resin in a ratio of 2:1 resin:ligand volume. The pH of the solution was adjusted to ~7-8 using 20x PBS and the mixture was rotated at 4°C overnight. The following day resin was pelleted (1400 rcf, 3 mins), supernatant discarded and then washed with 3x resin volumes of ice cold 0.1 M Tris-HCl [pH 8.5] via rotation for 3 hours at room temperature. After this the resin was pelleted (1400 rcf, 3 mins) and washed 3x with ice cold 0.1 M Tris-HCl [pH 8.5], 3x with ice cold 0.1 M sodium acetate, 0.5 M NaCl [pH 4.5], and with alternating washes (2x each) of ice cold 0.1 M Tris-HCl [pH 8.5] and ice cold 0.1 M sodium acetate, 0.5 M NaCl [pH 4.5]. Following washing, resin was stored at 4°C as a 33% solution in aqueous 20% EtOH.

In order to validate NAA50’s recognition properties, chemoproteomic capture and competition experiments were performed according to the previously reported method.^4^ Briefly, T-47D breast cancer proteomes in 1x PBS were diluted to 2 mg/mL in assay buffer (150 mM Tris-HCl [pH 8.0], 50 mM NaCl, 0.1% BSA). 30 uL of resin slurry (33% solution in 20% EtOH) was briefly washed once with 1 mL of ice-cold 1x PBS, combined with proteome (0.75 mg), and rotated 1 hour at 4°C. After incubation, the resin was pelleted (1400 rcf, 1:30 mins, 4°C) and the supernatant discarded. The resin was then subjected to 3x washes using ice cold wash buffer (50 mM Tris-HCl pH 7.5, 150 mM NaCl, 1.5 mM MgCl_2_, 5% glycerol). For each wash resin and wash buffer were rotated for 2 minutes at 4°C before the supernatant was discarded. Following the final wash, resin was collected on top of centrifugal filters (Pall, #ODM02C35). Proteins were eluted from the resin using 40 uL 1X SDS sample buffer. After the addition of SDS buffer, samples were boiled 10 mins at 95°C, then centrifuged (1400 rcf, 5 mins). This was performed 2x and the filtrates were combined, resulting in a final elution volume of 80 uL. Eluant was subjected to SDS-PAGE and western blotting analysis as described above (Materials and Methods). Chemoproteomic capture experiments using purified Nanoluc-NAA50 were carried out equivalently, replacing proteomes with 750 nM of purified recombinant protein in dilution buffer (150 mM Tris-HCl [pH 8.0], 50 mM NaCl, 0.1% BSA). Competition experiments incubated acetyl-CoA or CoA with proteome for 30 minutes on ice prior to addition of capture resin.

**BRET detection method**

For BRET experiments, Cy3-Ahx-CoA **1** (0-120 nM) was diluted in assay buffer (150 mM Tris-HCl [pH=8.0], 50 mM NaCl, 0.1% BSA) and added to Nanoluc-NAA50 (100 nM) in a 384-well plate. Plates were briefly centrifuged (500 rcf, 30s, room temperature) to remove air bubbles, covered to prevent photobleaching, and incubated at room temperature for 30 minutes. After incubation, furimazine was added to each well to a final concentration of 800 nM. Plates were briefly centrifuged once more, and immediately read using Cytation 5 plate reader. Spectral scans used a higher furimazine concentration (1600 nM) and were conducted over a range of 400-700 nm using a 5 nm step size. Endpoint readings were performed using a 460 nm band pass filter with a bandwidth of 40 nm (donor, Nanoluc) and a 590 nm band pass filter with a bandwidth of 35 nm (acceptor, Cy3) with an integration time of 1s. For ligand displacement assays, Cy3-CoA probe (**1** or **2**, 1 μM) was diluted in assay buffer (150 mM Tris-HCl [pH=8.0], 50 mM NaCl, 0.1% BSA) and added to Nanoluc-NAA50 (100 nM) in the presence of competitor metabolite. Acetyl-CoA and CoA competition was assessed over a concentration range of 10^-9^ M to 10^-3^ M (10-fold incremental steps), while propionyl-CoA, butyryl-CoA, malonyl-CoA, palmitoyl-CoA, and succinyl-CoA competition was assessed at a single concentration of 1 μM. Acyl-CoAs were prepared as aqueous solutions fresh prior to use with integrity verified by analytical LC-MS. The methods noted above were followed for plate preparation. All reactions were prepared and analyzed in ≥3 technical replicates unless otherwise noted. Values represent the average and error bars represent the standard deviation. Data were analyzed using Excel and GraphPad Prism 9 graphing software. Results are reported as a BRET ratio (Cy3/Nluc), which was calculated by dividing the raw emission value (RLU) measured for the Cy3 BRET acceptor (590/35 filter wavelength) by that of the Nanoluc BRET donor (460/40 filter wavelength):

*Cy3/Nluc BRET ratio =* $\frac{acceptor channel emission}{donor channel emission}$

For competition studies, results are represented as normalized emission compared to a control without competitor metabolite added:

*Normalized emission (Cy3/Nluc) =*

$(\frac{acceptor channel emission (+ metabolite)}{donor channel emission (+ metabolite)})\div(\frac{acceptor channel emission (- metabolite)}{donor channel emission (- metabolite)})\times100$

**BRET acetyltransferase assay**

Cy3-Ahx_4_-CoA **1** (1 μM) was diluted in assay buffer (150 mM Tris-HCl [pH=8.0], 50 mM NaCl, 0.1% BSA, 5 mM MgCl_2_, 1 mM DTT) and added to a pre-mixed solution of Nanoluc-NAA50 (100 nM), Histone H4 peptide [1-21] (50 uM), CPI-1612 (0-1 μM) and acetyl-CoA (1 μM) (Millipore Sigma, #A2181) in a 384-well plate. Furimazine was added to each well to a final concentration of 1 μM. Reactions were initiated by addition of EP300 at a final concentration of 75 nM. Plates were then centrifuged (500 rcf, 30s, room temperature) and immediately read using the Cytation5 plate reader. Endpoint emission measurements were taken every 2.4 minutes over an interval of 40 minutes total using a 460/40 band pass filter (Nluc donor) and a 590/35 band pass filter (Cy3, acceptor) with an integration time of 1s. The internal temperature of the plate reader was set to 37°C for the duration of the reaction. BRET ratios were calculated as described, with relative emission determined via normalization to a control without acetyl-CoA added. KAT6A (50 nM) was analyzed via a nearly identical protocol, substituting the appropriate enzyme and inhibitor (PF-9363), respectively, and increasing the furimazine concentration to 2 nM.

**Full gels and Western blot images**

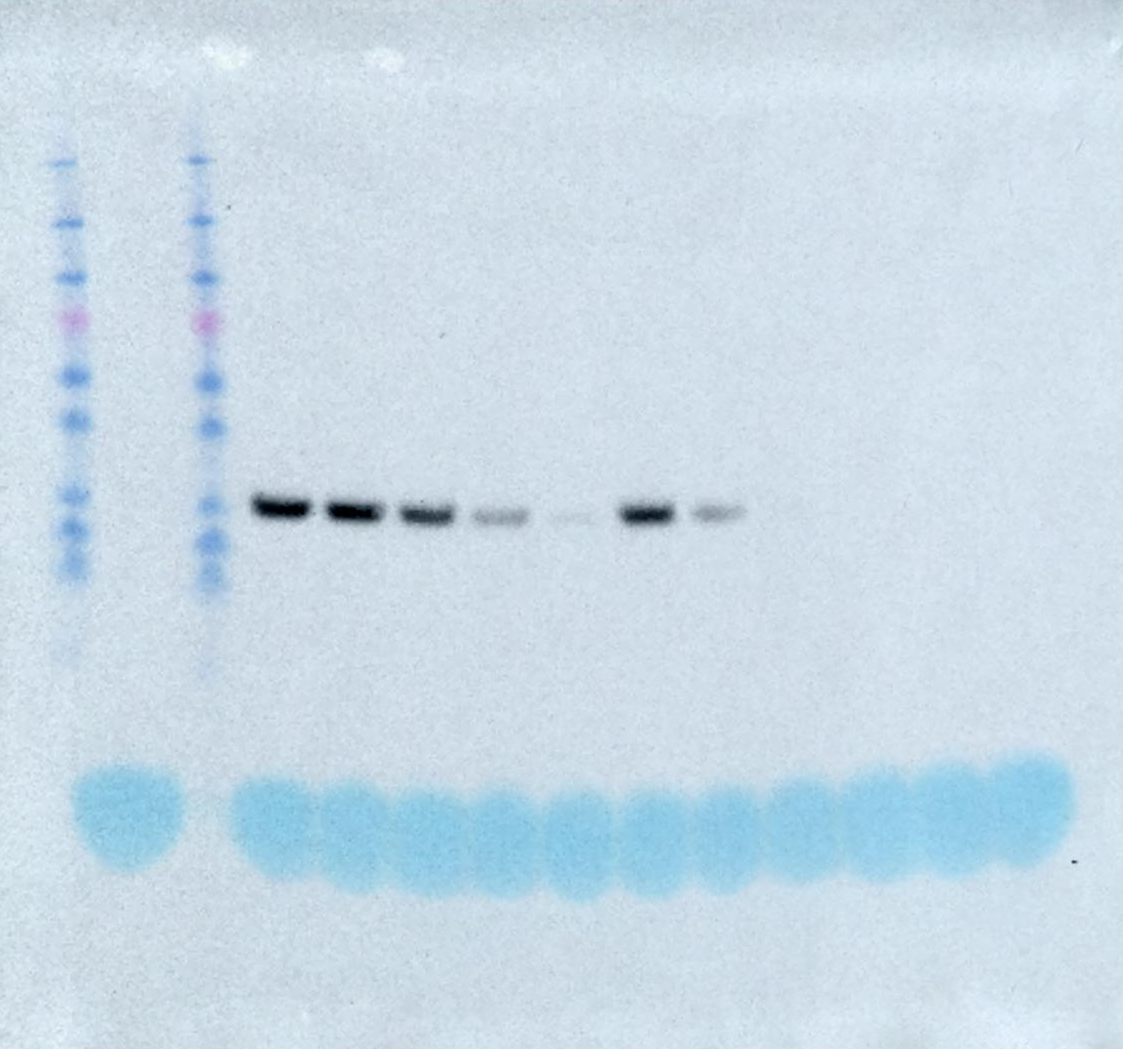

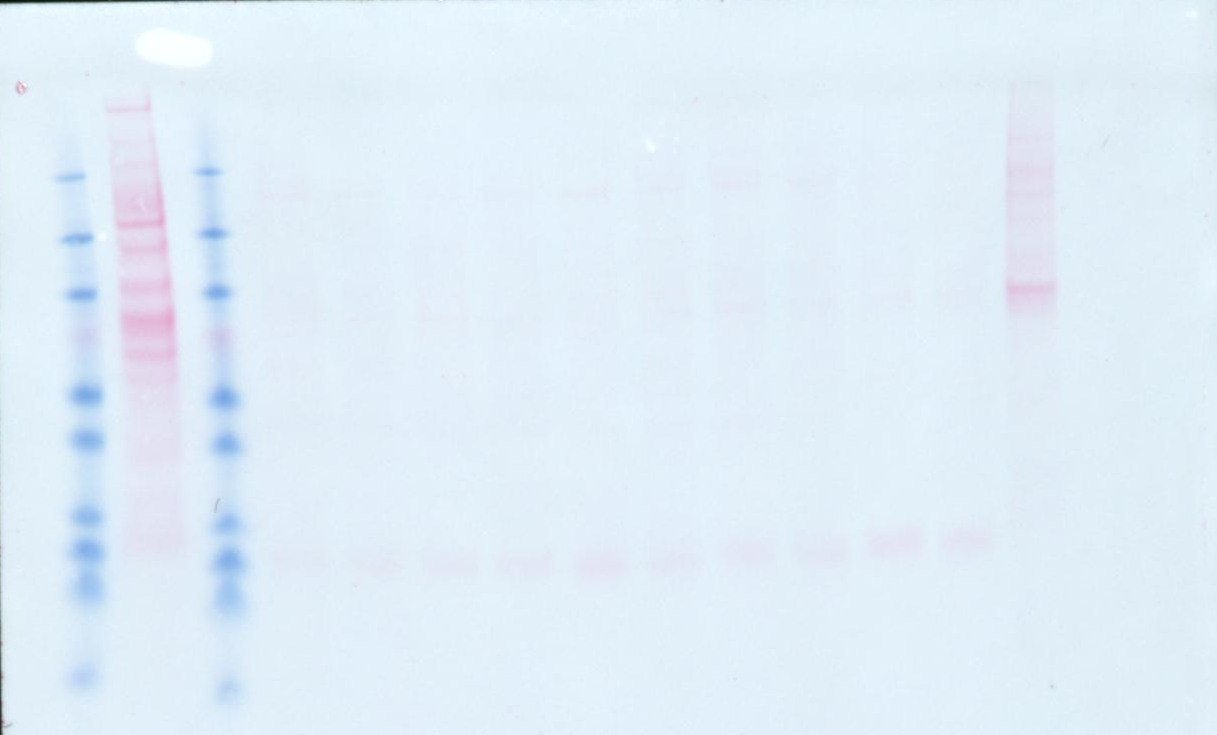

Full western blot for Figure 1. Top: anti-NAA50. Bottom: Ponceau. Note that Figure 1 contains a cropped version of this gel as CoA competition on left in native gel, but is on right in Figure 1.
